# Direct measurement of X-ray induced heating of microcrystals

**DOI:** 10.1101/603522

**Authors:** Anna J Warren, Danny Axford, Robin L Owen

## Abstract

Temperature control is a key aspect of macromolecular crystallography, with the technique of cryocooling routinely used to mitigate X-ray induced damage. Beam induced heating could cause the temperature of crystals to rise above the glass transition temperature, greatly increasing the rate of damage. X-ray induced heating of ruby crystals 20-40 microns in size has been quantified non-invasively by monitoring the emission wavelengths of X-ray induced fluorescence during exposure to the X-ray beam. For beamsizes and dose-rates typically used in macromolecular crystallography the temperature rises are of order 20 K. The temperature changes observed are compared with models in the literature and can be used as a validation tool for future models.

**Synopsis:** X-ray induced heating of micro-crystals is quantified through the temperature-dependent shift in X-ray induced fluorescence from ruby crystals.

## 1. Introduction

X-ray induced damage is an inevitable aspect of macromolecular crystallography at synchrotron sources (Garman, 2010). By far the most successful and widely implemented means of mitigating damage is to hold samples at ~100 K using an open flow nitrogen cryostat. A key tenet of this approach is the assumption that the crystal is held below the glass transition temperature of ~130-140 K (Johari *et al*., 1987, McMillan & Los, 1965, Sartor *et al*., 1994) during data collection (Weik *et al*., 2000). Above this temperature radical species are mobile within the crystal and, as a result, the rate at which radiation damage occurs greatly increases. During a diffraction experiment radical species are generated within the crystal through the X-ray induced radiolysis of water. This results in the formation of solvated electrons and hydroxyl radicals (Klassen, 1987). While electrons and holes are mobile down to 77 K (Jones *et al*., 1987, Symons, 1995), the mobility of hydroxyl radicals increases significantly between 115 K (Zakurdaeva *et al*., 2005) and 130 K (Symons, 1999). The energy deposited by the X-ray beam during data collection causes the temperature of the sample to rise above the nominal holding temperature of 100 K. As storage-rings and beamlines evolve, the increasing fluxes and flux densities available mean that this beam induced heating could cause the temperature of crystals to increase above the glass transition point. This would result in a greatly accelerated rate of damage and reduced crystal lifetimes. Beam induced heating is also a concern in room temperature macromolecular crystallography (MX) as reaction rates increase as temperature increases; a rule of thumb from the Arrhenius equation is that reaction rates double for every 10 K increase. Unless beam heating can be reduced, or outrun, any gains made by collecting data faster using more intense beams may be limited due to increased reaction and diffusion rates.

Despite the potentially deleterious implications of beam induced heating in synchrotron-based MX, a relatively limited amount of research has been carried out in this area. Helliwell (Helliwell, 1984) described a simple adiabatic model of beam induced heating. Later models take into account both conduction of heat within the sample and heat transfer away from the sample by the surrounding gas stream (Nicholson *et al*., 2001, Kriminski *et al*., 2003, Kuzay *et al*., 2001, Mhaisekar *et al*., 2005). In all cases it was shown that, as conductive heat transfer is more efficient that convection, temperature variations within the sample are expected to be small in comparison to the temperature difference between the sample and the surrounding gas stream.

In order to directly measure X-ray beam induced heating, an experimental approach was taken in which a thermal imaging camera was used to observe the heat rise within 1 mm and 2 mm diameter glass beads and validate theoretical predictions (Snell *et al*., 2007). In this study it was shown at a third generation synchrotron source that the temperature rise within a flow of nitrogen gas was of order 10-15 K: *i.e*. not enough to exceed the glass transition temperature. The continuous development of beamlines and synchrotron sources means however beamsizes are continually getting smaller with a concomitant increase in flux density. Furthermore, a 1mm glass bead has a volume some four or five orders of magnitude greater than a typical protein crystal which may be easily less than 50 microns in its longest dimension.

More recently (Mykhaylyk *et al*., 2017) reported non-contact luminescence lifetime cryothermometry for the *in-situ* measurement of different protein sample mounts within the vacuum vessel of beamline I23 at Diamond Light Source. A small piece of scintillating material (~100-200 µm) was used, where the luminescent decay time could be measured, and using calibration curves a temperature of the crystal could be derived. They were then able to test how effectively differing loop types kept the crystal cool at the sample position, observing a temperature range of 60-110 K, with the goniometer temperature held at 40 K. This method of temperature measurement proved interesting as the scintillating material has a volume more similar to that of a protein crystal, and like the IR experiments described above, the measurements were taken without the need for any physical connections to the crystal.

Here we report the investigation into the effect of heating on micron-sized crystals on the microfocus beamline I24 at Diamond Light Source. The (laser and X-ray) induced fluorescence of ruby crystals provides a convenient means of quantifying the X-ray induced temperature rise in small, protein crystal sized, samples. As with the Snell and Mykhaylyk experiments referenced above, this approach has the advantage that no cables or other physical connection between the sample and probe that might transfer heat to or from the sample are required. Ruby has the additional advantage that it is a well characterised system and temperature dependent changes in fluorescence have been the subject of many systematic studies. The most common means of inducing ruby fluorescence is with a green laser (Syassen, 2008), but X-ray induced fluorescence can also be exploited. In both cases, as the temperature of ruby increases the wavelength of fluorescence peaks also increases (Syassen, 2008, Ragan *et al*., 1992, McCumber & Sturge, 1963). For the experiments reported here this temperature dependent line shift provides a convenient micron sized temperature probe.

Experimental measurement of beam induced heating in ruby crystals allows validation of beam heating models and hence prediction of whether new sources and beamlines will cause the temperature of protein crystals to increase beyond tolerable limits.

## 2. Materials and Methods

### 2.1. Beamline / spectrometer parameters

Data were collected at beamline I24 at Diamond Light Source using a beamsize of 20 × 20 µm^2^ (dimensions are FWHM; Gaussian profile, uncollimated beam) and X-rays of energy 12.8 keV, with a flux of 1.19 × 10^12^ ph s^−1^, and 9.2 keV with an incident flux of 3.18 × 10^12^ ph s^−1^. Photon fluxes were measured using a silicon PIN diode (Owen *et al*., 2009). These flux densities allowed dose-rates of up to 1.5 MGy s^−1^ (12.8 keV) and 7.4 MGy s^−1^ (9.2 keV) to be realised in ruby samples. Absorbed doses were calculated using RADDOSE-3D (Zeldin et al., 2013). The dose values quoted and used below are the average diffraction weighted dose reported by this program. In RADDOSE-3D the small molecule option was utilised allowing zero solvent content to be specified.

Emission spectra were collected using mirrored lenses (Bruker) mounted in an off-axis geometry (Figure 1a). The lenses were positioned so the focus of each coincided with the intersection of the goniometer rotation axis and X-ray beam (Figure 1b). Absorption was monitored over the 560-830 nm wavelength range using a Shamrock 303 imaging spectrograph (Andor). The spectrograph was calibrated to better than 0.01 nm using a mercury argon lamp (Ocean Optics) and the calibration confirmed by cross comparison of spectra collected on a second spectrograph with a fixed grating.

**Figure 1.**
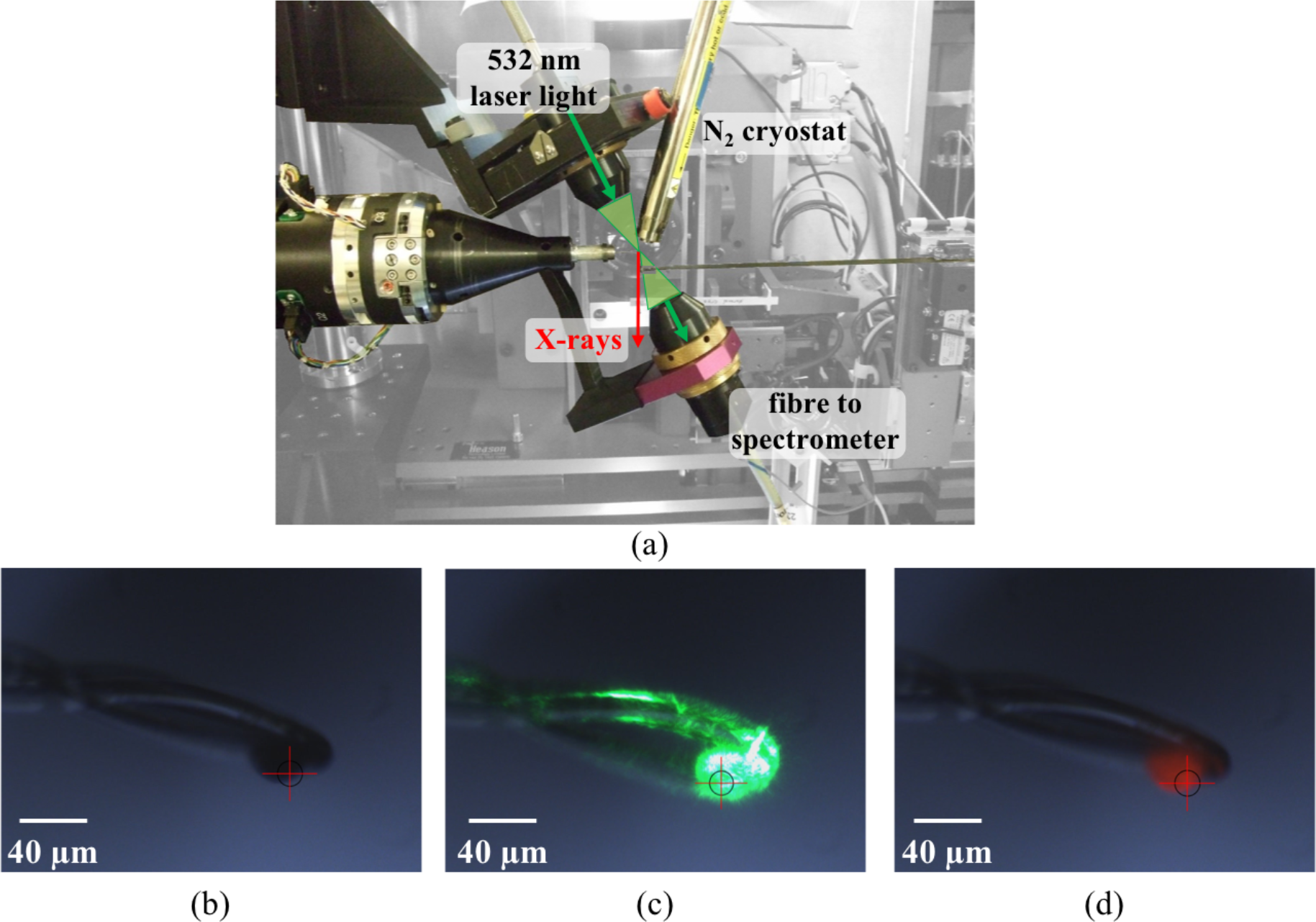
(a) Experimental set up on beamline I24 showing the sample position with the spectrometer installed and the cryostream in place to control the temperature of the experiments. (b-d) View of a 40 µm ruby sphere mounted (b) within a nylon loop at the sample position, (c) illuminated with the 532 nm laser and (d) when exposed to the X-ray beam, demonstrating the fluorescence of the ruby.

For laser-induced fluorescence a 532 nm laser was used for excitation (shown in Figure 1c). The laser power at the sample position was 13 µW (measured using a Thorlabs power meter). Each measurement was taken for 0.2 s with an accumulation of five images, except where time evolution studies were carried out in which 5 ms, 10 ms and 20 ms images (1 accumulation) were used. In the setup described, X-ray induced fluorescence (shown in figure 1d) was an order of magnitude more intense than that induced by the laser (Figure 2a).

**Figure 2.**
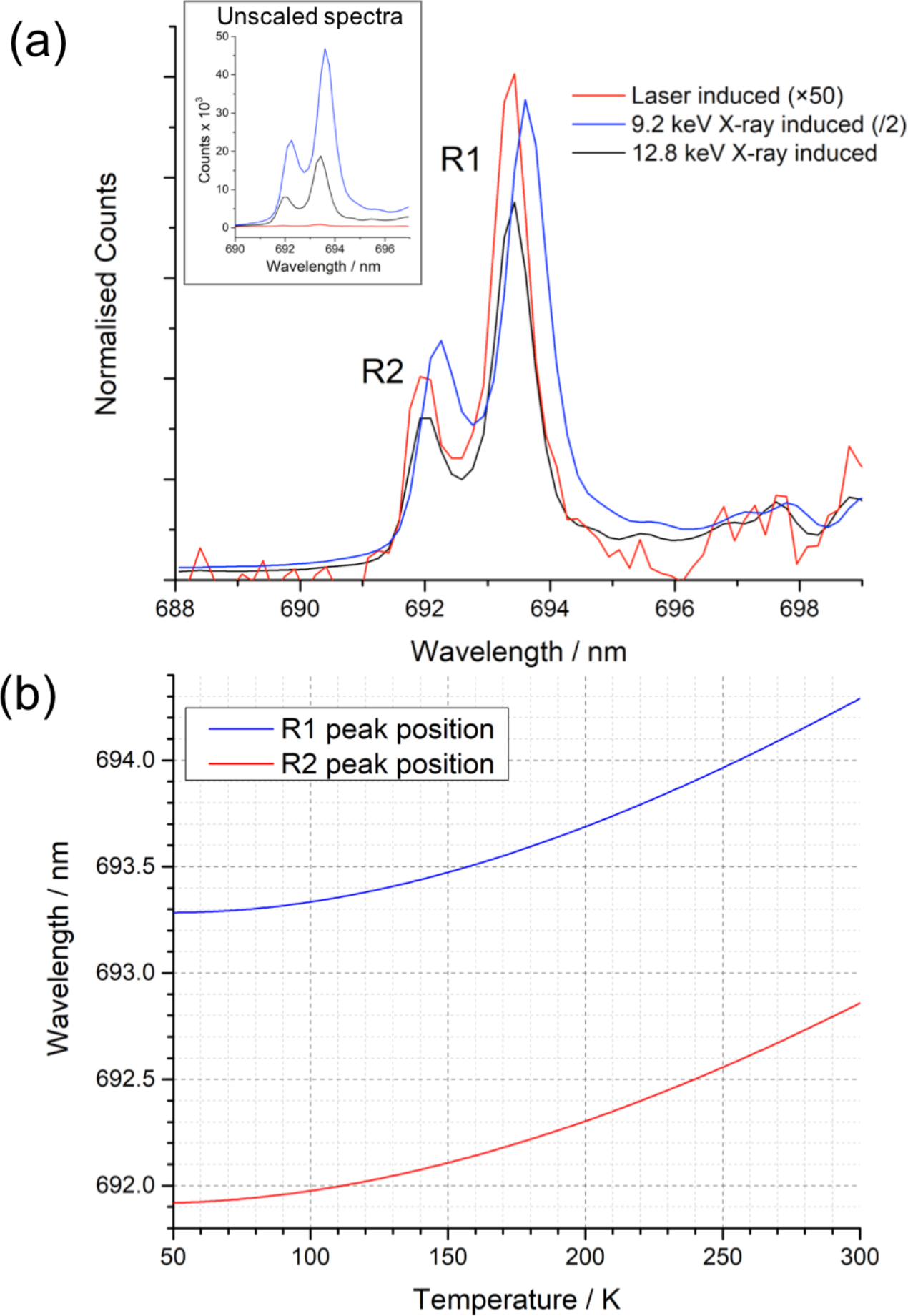
(a) Laser and X-ray induced fluorescence of ruby showing peak shifts and relative intensity of laser and X-ray induced spectra. In addition to being red-shifted, the R1 and R2 peaks become broader at higher temperatures. The inset shows the unscaled spectra. (b) Calibration curve showing the position in wavelength of the ruby R1 and R2 fluorescence peaks obtained from equations 2 and 3. Note that the cubic function has a stationary point at 50 K (R1) and 40 K (R2), so temperatures below this would result in a peak with the same wavelength at a higher temperature. Temperatures below 80 K were not investigated as part of this study.

### 2.2. Sample Preparation

Ruby spheres were purchased from diamondANVILS^1^ with diameters varying between 10 – 50 µm. Spheres were mounted on nylon loops using as little Fomblin Y oil (Sigma-Aldrich) as possible and placed directly on the beamline and cooled in an open flow nitrogen cryostat held at 100 K. Ruby is aluminium oxide doped with a small amount of chromium: the asymmetric unit of ruby is (Al_0.33_Cr_0.00333_)O_0.5_ in space group *R*-3*c* and cell parameters *a* = 4.75Å *c* = 12.99 Å. The aluminium and chromium share a third occupancy on a special position, and the oxygen has half occupancy also on a special position.

For all data collected the nitrogen cryostat was set to the default flow rate of 10 l min^−1^. For experiments using a helium cryostat it was not possible to set the helium flow rate to a specific value. The flow was set so it was approximately the same as that of nitrogen based on the helium consumption of the cryostat given by the manufacturer ^2^, the gas expansion coefficient of helium, and visually keeping the flow rate constant over the duration of the experiment.

### 2.3. Data analysis

Laser induced ruby R1 and R2 fluorescence peaks are shown in Figure 2a and are best described by a Lorentzian profile (Ragan *et al*., 1992). OriginPRO was used to fit the experimental data with a double Lorentzian of the form

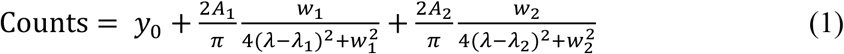

where *y*_0_ an offset and *A*_*i*_ is the area, *w*_*i*_ the width, and *λ*_i_ the centre of each peak. The wavelength dependence of the R1 and R2 fluorescence peaks as a function of temperature, T, was determined from the functions given by (Ragan *et al*., 1992). The peak positions in cm^−1^ are given by

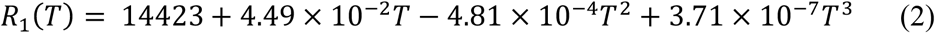

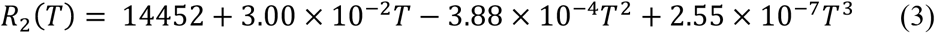

Calibration curves relating wavelength (1/wavenumber) to temperature derived from equations 2 and 3 are shown in Figure 2b. As R1 is the larger peak, the position of this peak was used to calculate the temperature of the crystal. The difference in temperature calculated using the position of R1 versus that calculated from the position of R2 was typically less than 3 K. For brevity we refer to the temperature measured and calculated in this way as the ‘fluorescence temperature’ below, and all ruby crystal temperatures refer to the observed fluorescence temperature.

## 3. Results

Emission spectra were collected from two ruby crystals, the first with a diameter of 20 µm (*i.e*. matching the beamsize) and the second with a diameter of 40 µm. Laser induced emission spectra were relatively weak (Figure 2a), and data collection at different laser powers resulted in no discernible peak shift. Crystal temperatures recorded using the laser alone were therefore taken to be representative of the temperature of ruby crystals held in the nitrogen flow in the absence of external heating. In the absence of X-rays, the difference in fluorescence temperature and nominal set point of the nitrogen cryostat was ~15 K with the cryostat set to 100 K. Note that this refers to the fluorescence temperature determined from optical laser induced fluorescence when probe induced heating of the sample is expected to be almost zero. The difference in fluorescence temperature and the cryostat setpoint was observed to decrease to zero as the cryostat setpoint temperature increased from 100 K to room temperature (Figure 3). This difference may originate from an error in the cryostat calibration or imperfect positioning of the sample in the gas stream. All subsequent data refer to beam induced temperature changes and so are unaffected by this offset.

**Figure 3.**
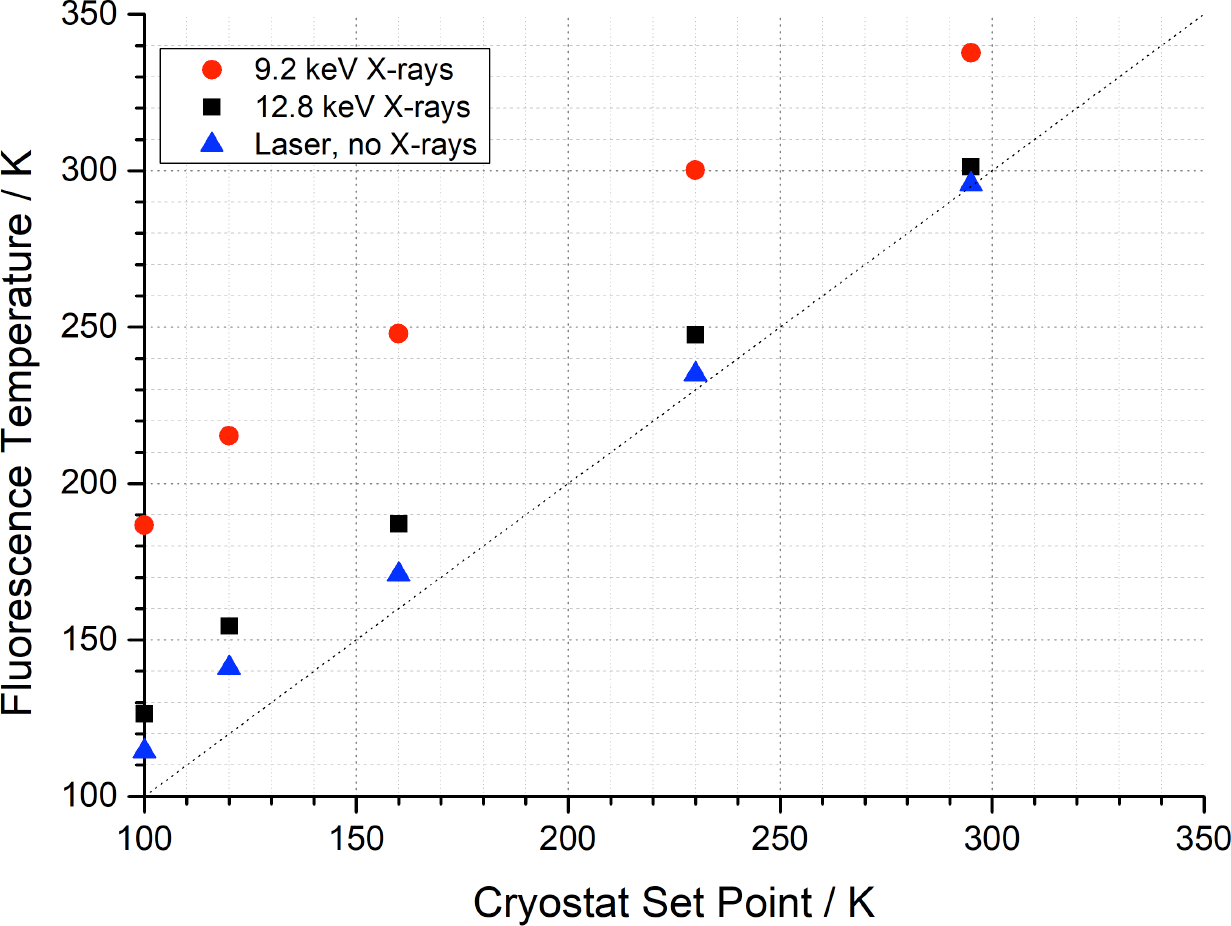
Optical laser (no beam) and X-ray fluorescence temperature of a 40 µm ruby crystal as a function of cryostat set point. The X-ray beamsize was uncollimated 20 × 20 µm^2^ Gaussian and incident flux 3.18 × 10^12^ ph s^−1^ and 1.19 × 10^12^ ph s^−1^ at 9.2 keV and 12.8 keV respectively (detailed in §2.1).

The observed change in temperature as a function of dose-rate is shown in Figure 4. It can be seen that the rate of change is independent of energy for a given crystal size and is a function of absorbed dose. The large X-ray cross-section of ruby means that extremely high dose-rates are reached at both X-ray energies. The dose-rates at 12.8 keV are more representative of those realised in protein crystallography (< 5 MGy s^−1^). At this energy, for the 40 µm ruby crystal the maximum increase in temperature is 15 K, while for the 20 µm crystal the maximum temperature rise is 5 K. Linear fits to the data are overlaid, and the intercepts (i.e. temperature rise induced at a dose-rate of 0 Gy s^−1^) are 1.3 ± 0.8 K (20 µm crystal) 3.5 ± 0.6 K (40 µm crystal). The deviation of these (and intercepts in figure 5) from 0 K reflects the uncertainty in the measurements and could arise from experimental error in several parameters such as X-ray flux and beamsize, crystal size and fluorescence wavelength.

**Figure 4.**
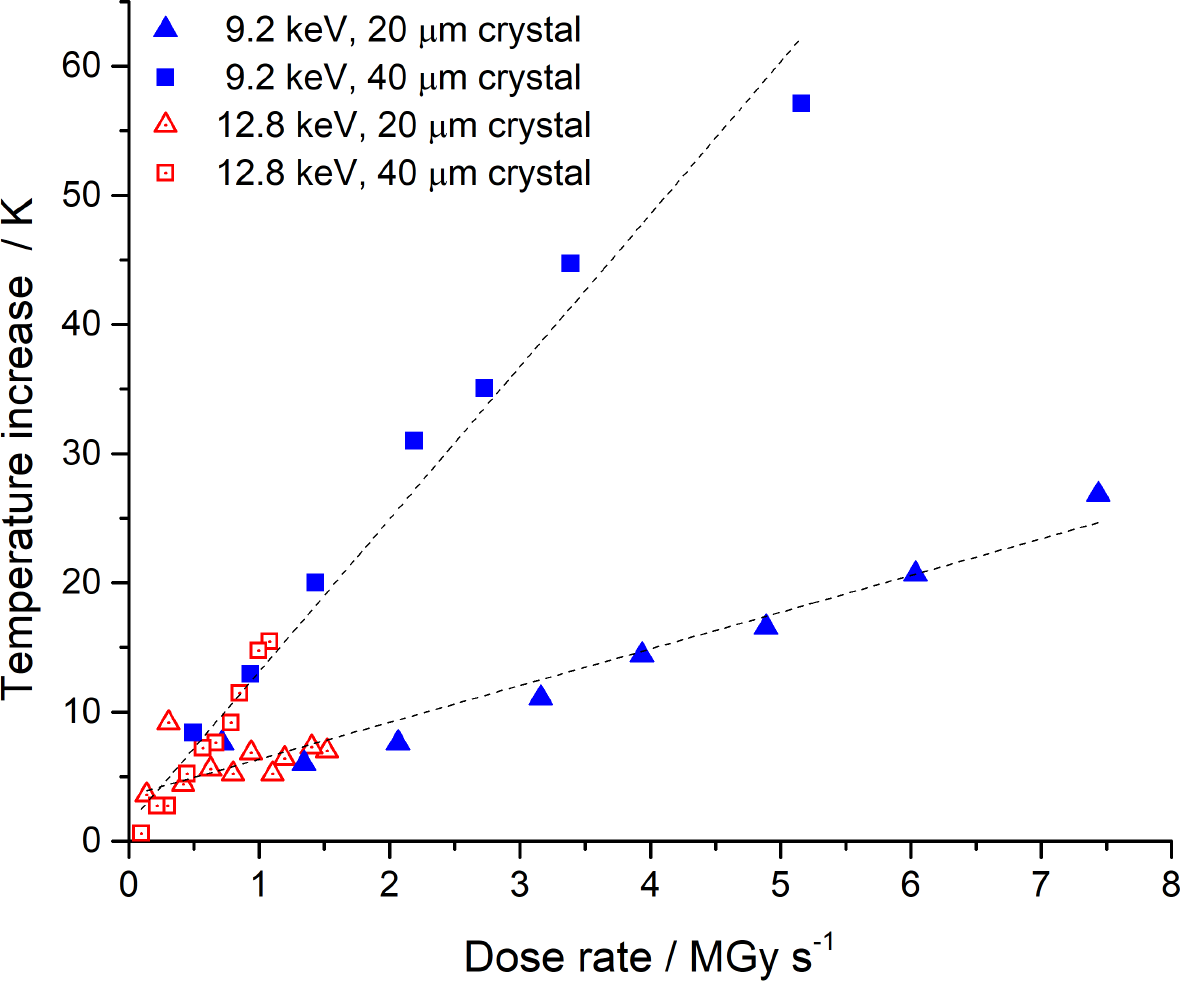
The effect of dose rate on the temperature increase of ruby crystals. Blue and red points indicate data collected at 9.2 and 12.8 keV respectively. Data collected with the crystal size matched to the beam are shown as triangles, squares indicate the crystal was larger than the beam.

**Figure 5.**
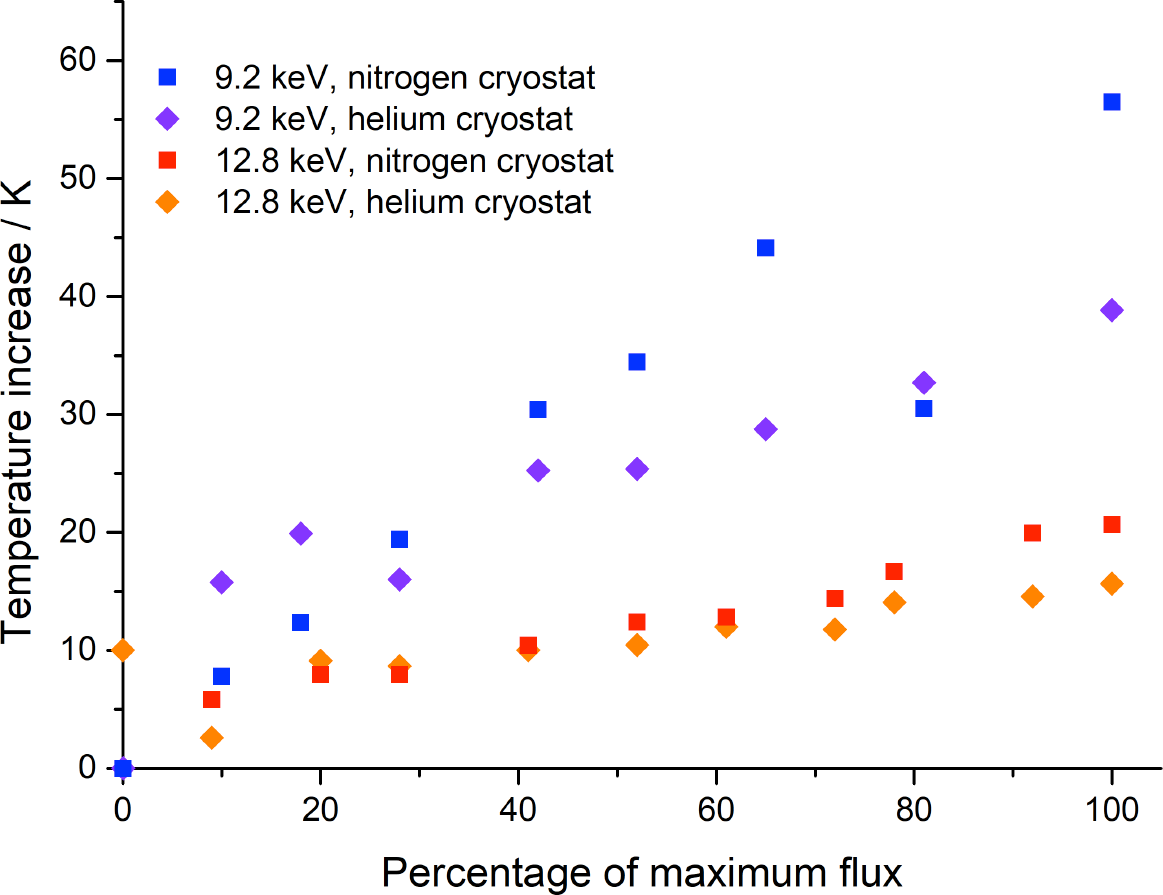
Effect of cryostat gas on change in temperature. To allow direct comparison of data collected at 9.2 and 12.8 keV, temperature changes are plotted as a percentage of maximum flux rather than flux density or dose. The flux densities and resulting absorbed doses are the same as those shown in Figure 4. Data collected from a 40 µm diameter ruby sphere.

The effect of the gas used to cool the sample is shown in Figure 5. To allow direct comparison of data collected at 9.2 and 12.8 keV, data are plotted as a function of the maximum flux as shown in Figure 4. The use of helium reduces the X-ray induced temperature rise. In the case of 9.2 keV X-rays the temperature increase at maximum flux is reduced from 58 K to 39 K, while the change is reduced from 21 K to 15 K for 12.8 keV X-rays: a reduction of ~30% in each case. The use of helium as a cryogen clearly significantly reduces X-ray induced temperature increases and this could be particularly useful when extremely brilliant sources are used and samples are subject to extremely high dose-rates. However any gains in keeping crystals below the glass transition temperature have to be tensioned against the ease of use and ubiquity of nitrogen cryostats, the finite availability and cost of helium, and the extremely short lifetime of protein crystals in such intense X-ray beams.

In order to quantify the time required for samples to reach a steady-state temperature in the X-ray beam, the wavelength shift of ruby fluorescence was recorded as a function of time (Figure 6). It can be seen that the initial rate of change of temperature is extremely large (> 6000 K s^−1^), but a steady-state fluorescence temperature is reached relatively quickly after only ~30 ms (~150 kGy). An exponential fit of the form Temperature rise = T_0_ + A_0_ exp(−dose/d_0_) where T_0_, A_0_ and d_0_ are constants was fitted to data with the shortest integration time (5ms). In this case, the quantity d_0_ was 49 ± 5 kGy meaning the dose required for half of the final temperature rise to be reached was just 34 kGy (~6 ms).

**Figure 6.**
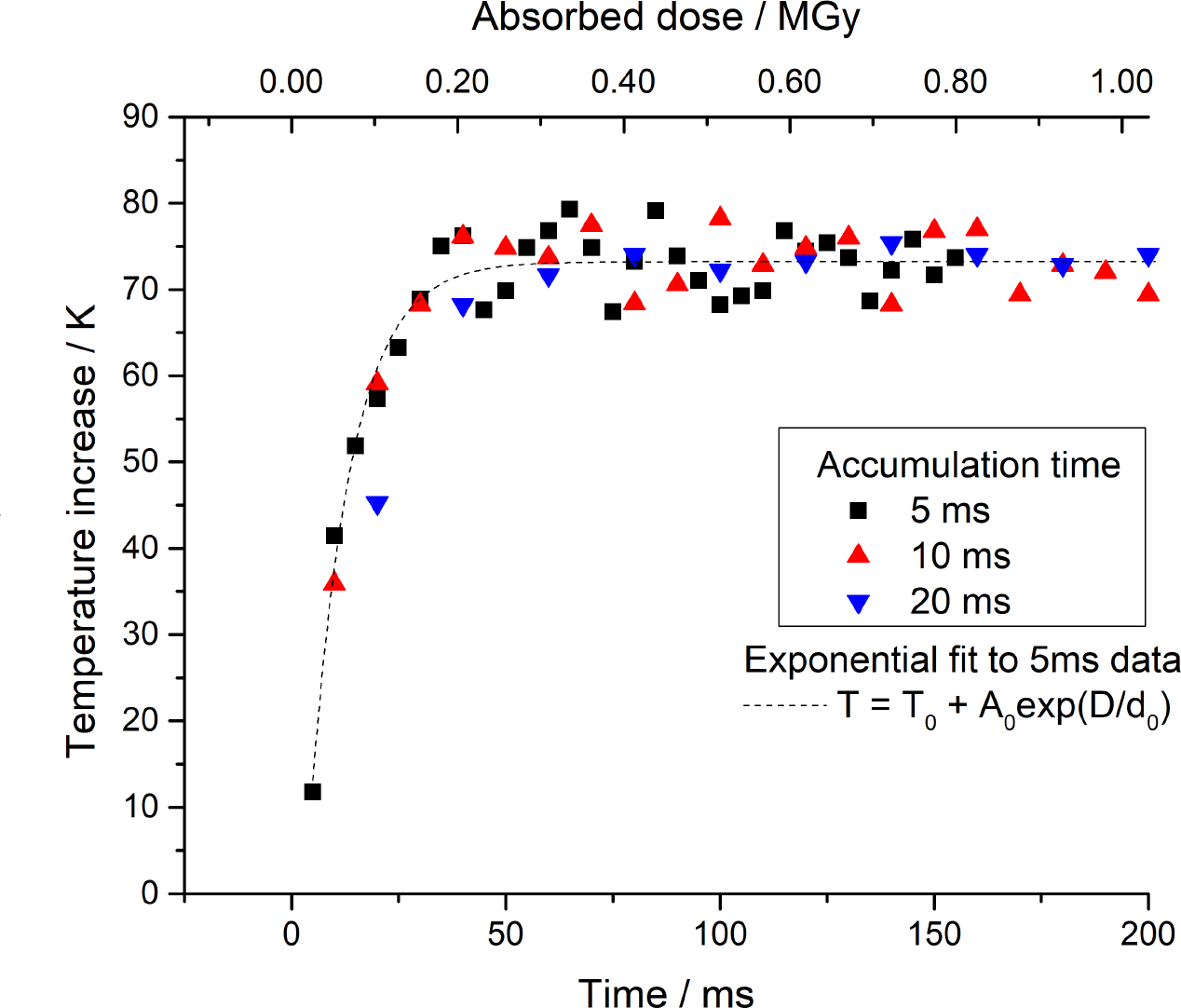
Rate of change of fluorescence temperature induced by 9.2 keV X-rays in a 40 µm ruby crystal. Data were collected with a 5 ms, 10 ms and 20 ms accumulation time: 5 ms data points exhibit more noise but greater temporal resolution. Initial rate of change of temperature (over first 10 ms) is > 6000 K s^−1^, with a steady state temperature is reached after ~ 40 ms. Data collected at an incident dose rate of 5.1 MGy s^−1^.

## 4. Discussion and Conclusions

Several models for X-ray induced heating of samples have been proposed, we here briefly summarise them and compare the calculated temperature rises of ruby with those experimentally observed. The most basic model of beam induced temperature changes is an adiabatic one *i.e*. ignoring any heat exchange. In this case the temperature rise can be calculated by dividing the energy absorbed by the mass and the specific heat capacity of the sample (symbols used in the following equations are defined in Table 1). As the dose is the energy absorbed per unit mass *Q/m* can be replaced by the absorbed dose D

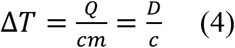

**Table 1.**
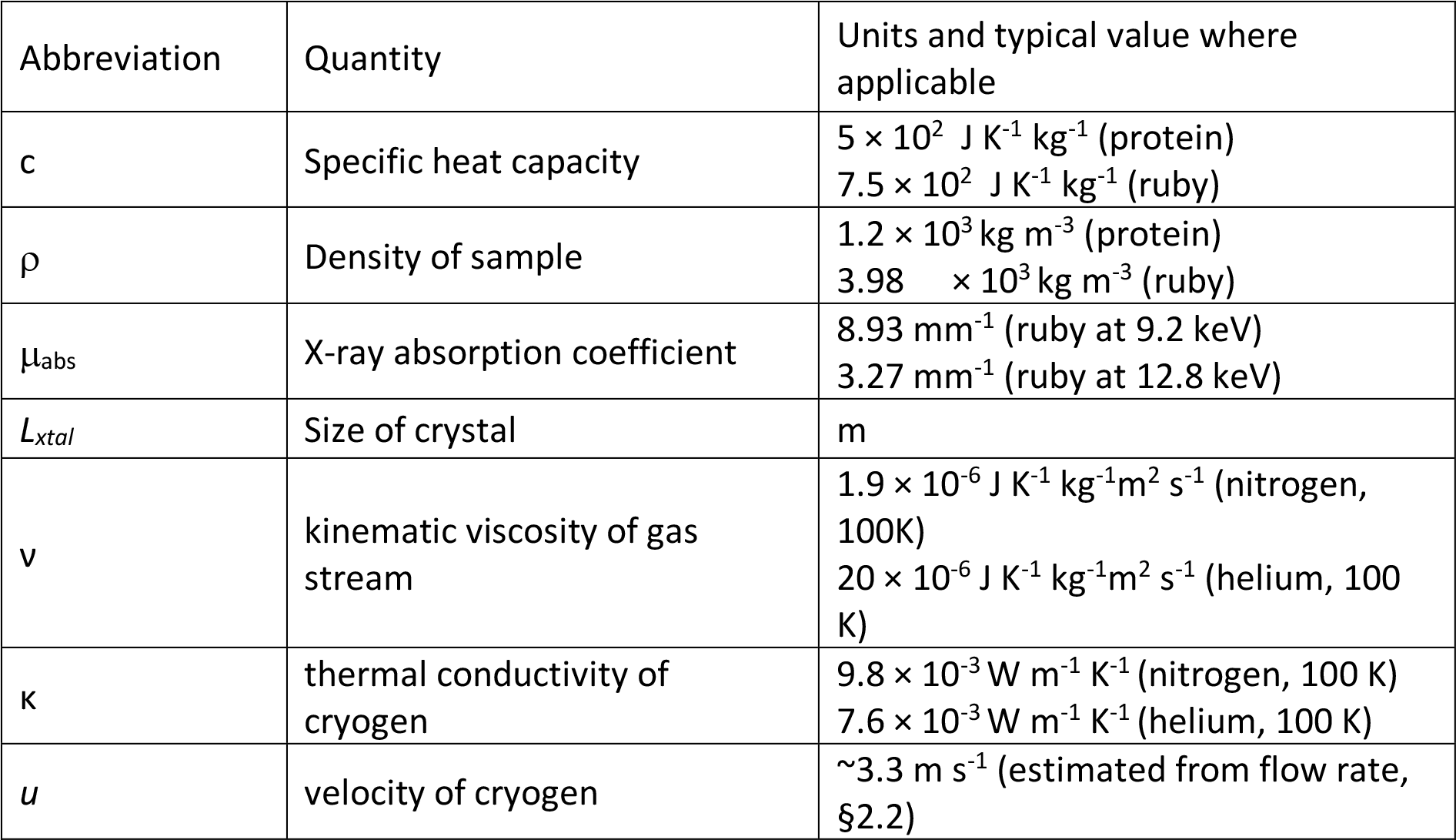
Physical parameters relevant to sample X-ray beam heating and their abbreviations and values as used in this work. Kinematic viscosities and thermal conductivities were taken from (Kriminski *et al*., 2003).

| Abbreviation | Quantity | Units and typical value where applicable |
| --- | --- | --- |
| $c$ | Specific heat capacity | $5 \times 10^2 \text{ J K}^{-1} \text{ kg}^{-1}$ (protein)<br>$7.5 \times 10^2 \text{ J K}^{-1} \text{ kg}^{-1}$ (ruby) |
| $\rho$ | Density of sample | $1.2 \times 10^3 \text{ kg m}^{-3}$ (protein)<br>$3.98 \times 10^3 \text{ kg m}^{-3}$ (ruby) |
| $\mu_{\text{abs}}$ | X-ray absorption coefficient | $8.93 \text{ mm}^{-1}$ (ruby at 9.2 keV)<br>$3.27 \text{ mm}^{-1}$ (ruby at 12.8 keV) |
| $L_{\text{xtal}}$ | Size of crystal | m |
| $\nu$ | kinematic viscosity of gas stream | $1.9 \times 10^{-6} \text{ J K}^{-1} \text{ kg}^{-1} \text{ m}^2 \text{ s}^{-1}$ (nitrogen, 100K)<br>$20 \times 10^{-6} \text{ J K}^{-1} \text{ kg}^{-1} \text{ m}^2 \text{ s}^{-1}$ (helium, 100 K) |
| $\kappa$ | thermal conductivity of cryogen | $9.8 \times 10^{-3} \text{ W m}^{-1} \text{ K}^{-1}$ (nitrogen, 100 K)<br>$7.6 \times 10^{-3} \text{ W m}^{-1} \text{ K}^{-1}$ (helium, 100 K) |
| $u$ | velocity of cryogen | $\sim 3.3 \text{ m s}^{-1}$ (estimated from flow rate, §2.2) |

We refer to this as the ‘basic model’. Temperature rises for the samples and beam parameters used in this work have been calculated and are shown in Table 2. It can be seen from the table that the basic model poorly describes heating of crystals by X-rays with temperature rises of hundreds of Kelvin predicted in tens of milliseconds, reflecting the limitations of such a simple model.

**Table 2.** Comparison of predicted and observed X-ray induced beam heating. The models used are described in the main text. The temperature change calculated using the basic model assumes 30 ms of exposure to X-rays (the approximate time taken to reach a steady-state temperature Figure 6). The KKT model and observed temperature rises are steady-state temperatures. The dose-rate is calculated from the average diffraction weighted dose reported by RADDOSE-3D.

| Sample | Energy / keV | Sample size / $\mu\text{m}$ | Cryogen | Dose rate / $\text{MGy s}^{-1}$ | $\Delta T$ / K (Basic model) | $\Delta T$ / K (KKT model) | $\Delta T$ / K Observed |
| --- | --- | --- | --- | --- | --- | --- | --- |
| Ruby | 9.2 | 20 | Nitrogen | 7.44 | 296 | 61.6 | 27 |
|  |  | 40 | Nitrogen | 5.16 | 205 | 120.6 | 58 |
|  |  |  | Helium | 5.16 | 205 | 50.5 | 39 |
|  | 12.8 | 20 | Nitrogen | 1.52 | 60 | 12.5 | 7 |
|  |  | 40 | Nitrogen | 1.08 | 43 | 25.2 | 21 |
|  |  |  | Helium | 1.08 | 43 | 10.5 | 15 |
| Protein | 9.2 | 20 | Nitrogen | 2.30 | 138 | 5.7 | - |
|  |  | 40 | Nitrogen | 1.68 | 101 | 11.8 | - |
|  |  |  | Helium | 1.68 | 101 | 5.0 | - |
|  | 12.8 | 20 | Nitrogen | 0.458 | 27 | 1.4 | - |
|  |  | 40 | Nitrogen | 0.336 | 138 | 2.4 | - |
|  |  |  | Helium | 0.336 | 138 | 1.0 | - |

(Kuzay *et al*., 2001) demonstrated that for all but the briefest time periods it is necessary to consider the convection of heat away from the sample when determining the temperature change induced by X-rays: crystals are heated internally by X-rays and externally cooled at the surface by the gas stream. This was also taken into account by Nicholson *et al*. (2001), though the Finite Element Analysis model described was not available to us and so is not compared with the data collected here. The Kuzay model of X-ray induced temperature change within a sample was refined by Kriminski, Kazmierczak and Thorne (Kriminski *et al*., 2003) (KKT) who showed that the temperature change at the surface of a sample relative to that of the bulk gas stream surrounding it is given by

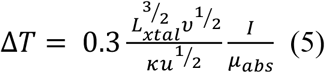

The assumption made that the absorbed power *p*_*abs*_ ≈ *IV*/*μ*_*abs*_, where *V* is the volume of the sample, can be used to reformulate this in terms of the more familiar absorbed energy per unit mass or dose, *D*, rather than the X-ray energy flux per m^2^.

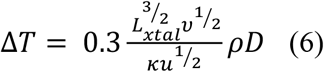

Again, symbols used are defined in Table 1. We refer to equation 6 as the KKT model below. It should be noted that in this equation the absorbed dose D is the absorbed dose per second, or dose-rate, rather than the total dose absorbed over the duration of the experiment. It can be seen from Table 2 that the temperature rises predicted by the KKT model agree well with the fluorescence temperature of the ruby crystals. Both predicted and observed temperature rises follow the same trends, with larger crystals showing larger temperature rises and a helium cryostat providing more efficient cooling. Temperature increases are over-estimated by a factor of ~2 in the case of ruby crystals in a nitrogen stream irradiated at 9.2 keV but are somewhat closer (within ~6 K) in all other cases. The reason for the discrepancy is not clear but may result from underestimation of the nitrogen flow or overestimation of the crystal size in these experiments. As an example, if the size of the 40 micron crystal was overestimated by 5 microns, then the predicted steady-state temperature increase would reduce from 121 K to 101 K.

Heat transfer within a crystal is more efficient than heat transfer through the surrounding gas stream. It might therefore be expected that increasing the size of the crystal beyond that of the X-ray beam would result in a ‘fin effect’ where unirradiated regions of the crystal act as a heat sink. In this case the temperature rise for cube-shaped crystals would be reduced by a factor of (L_beam_/L_crystal_)^½^ (Kriminski *et al*., 2003). The absence of this effect can be accounted for by the Gaussian profile of the beam: there is significant X-ray intensity in the tails of the beam, beyond the full width half maximum quoted beamsize, resulting in the observed and predicted larger temperature rise in 40 micron ruby crystals. The beam intensity also varies along the beam path through the crystal. The X-ray attenuation lengths in ruby at 9.2 keV and 12.8 keV are ~110 µm and 290 µm respectively (calculated using *RADDOSE-3D*). Thus, in the case of a 40 micron ruby crystal the beam intensity falls by 30% (9.2 keV X-rays) or 13% (12.8 keV X-rays) which may result in over-estimation of modelled temperature rises. The (Snell *et al*., 2007) study was unique in that it was able to resolve temperature changes both temporally and spatially. In cases such as this, when the beam intensity varies significantly across the sample and also when the sample size is significantly larger than the X-ray beam (as was the case with the 1 and 2 mm glass beads used by Snell and coworkers) significant steady-state temperature gradients may result within the sample. and ‘single temperature methods’ such as X-ray induced fluorescence can offer only a partial view of beam induced heating.

The agreement between the predicted and observed temperature changes in ruby gives confidence in the predicted temperature increases in protein crystals. The temperature increases predicted (Table 2) at 12.8 keV are relatively small (< 3 K). It should be noted however that the beamsizes and fluxes used in this study are (now) relatively modest: a three-fold increase in flux has recently been realised at I24, while the beam can also be more tightly focused. If a protein crystal matched to a tophat beam of 5 × 5 µm^2^ is exposed to 3 × 10^12^ ph s^−1^ 12.8 keV X-rays the predicted temperature rise increases threefold to 9 K (dose rate 28 MGy s^−1^). While this should still result in a protein crystal remaining below the glass transition temperature, if heavy atoms such as selenium are added to the crystal composition then the absorbed dose increases significantly as does the predicted temperature rise to 26 K. Decreasing the energy of the incident X-rays means higher dose-rates are more easily realised with a concomitant increase in beam induced heating. As the X-ray induced temperature change in crystals is proportional to the absorbed dose (KKT model described above), this may provide motivation for data collection at higher energies when extremely intense X-ray beams are used. For small samples this gain may be further increased by the reduction in absorbed dose resulting from photoelectron escape (Cowan & Nave, 2008). In Figure 3 an offset of ~15 K between the fluorescence temperature and nominal set point of the nitrogen cryostat was observed. If a temperature offset due to inefficient cryocooling or alignment of a cryostat is combined with beam induced heating then it is clear that increased rates of damage may result and crystal lifetimes in conventional nitrogen cooled cryocrystallography may be significantly less than expected.

In conclusion, we have shown that ruby crystals and their fluorescence can be used as a convenient method for measuring the X-ray induced temperature changes. These results correlate well with the KKT model. With the beam parameters used here this model predicts X-ray induced temperature rises of < 3K in protein crystals with the result that they should remain well below the glass transition temperature. At current and near-future synchrotron beamlines significantly higher flux densities can be achieved however, and the heating of samples to temperatures in excess of this remains a concern and challenge for microfocus MX.

1 https://www.diamondanvils.com/

2 Cryo Industries of America

